# Decoding hypnotic consciousness: neural and experiential insights into induced and ideomotor suggestions

**DOI:** 10.1101/2025.05.11.652707

**Authors:** Juliette Gelebart, Alexandre Fouré, Romain Quentin, Ursula Debarnot

## Abstract

Hypnotic induction and ideomotor suggestions provide a powerful framework for investigating the remarkable capacity of verbal influence to reshape conscious experience, cognition, and motor control. We employed a multimodal approach combining high-density EEG, respiratory and behavioral monitoring, and first-person reports across three conditions: baseline resting state, progressive hypnotic induction (Light and Deep states), and an ideomotor task comparing a hypnotically suggested arm catalepsy to a voluntary simulation. EEG results revealed that light hypnosis was associated with early parieto-occipital alpha suppression and increased theta-band activity. As hypnosis deepened, frontoparietal connectivity increased in the theta while parasympathetic activation declined, challenging the view of hypnosis as a passive, low-arousal state and instead pointing to active top-down reorganization of large-scale brain networks. During the ideomotor state, participants exhibited distinct patterns of behavioral responsiveness, classified as tremblers and non-tremblers, despite reporting comparable disruptions in the sense of agency. Phenomenological analyses corroborated these distinctions, revealing that Tremblers attempted to move despite experiencing the action as involuntary or constrained, whereas Non-Tremblers refrained from acting due to a perceived impossibility or an inability to initiate the motor command. EEG connectivity analysis in Tremblers showed an increased frontoparietal gamma activity and reduced delta connectivity, suggesting heightened sensorimotor integration and greater executive monitoring under motor conflict. Together, these findings demonstrate that hypnosis engages dynamic top-down processes that reconfigure both neural connectivity and subjective experience depending on suggestion-types. They support predictive coding accounts of agency disruption and underscore the value of neurophenomenological methods for advancing consciousness science and informing clinical applications.

---

Hypnosis is increasingly understood as a top-down regulatory process where verbal suggestions can elicit mental representations that dynamically influence perceptual, cognitive, and ideomotor processes, leading to measurable changes in subjective experience and behavior (1, 2). In typical hypnosis session, three overlapping stages are driven by different suggestion-types (3): an induction state, where suggestions capture attention, enhance absorption in inner experiences, and reinforce hypnosis-related expectancies (4); a suggestion state, where ideomotor-like suggestions modulate motor control and the sense of agency, a key component of intentionality (5); and a de-induction in which suggestions reorient attention and restores ordinary perception and behavior. Underlying these stages, the top-down regulation of suggestions in hypnosis is a multifaceted process that integrates cognitive mechanisms such as attention, executive control, and monitoring, all shaped by sociocultural and psychological factors (6, 7). This complexity gives rise to inter-individual variability in hypnotic responsiveness (1, 8), as empirical evidence indicates that individuals may engage diverse cognitive strategies in response to similar suggestions, which complicates the isolation of suggestion-specific responsiveness across subjective, behavioral, and neural levels (9, 10). To date, and despite advances in the cognitive neuroscience of hypnosis, the neurophysiological and subjective responsiveness underlying induction and ideomotor suggestions, the most commonly used in hypnosis, remain largely elusive (1, 6). Yet, understanding these mechanisms could offer insights into broader theories of consciousness dynamics and inform the clinical application of hypnosis-based interventions (11, 12).

Advancing the understanding of induction and ideomotor suggestions requires addressing methodological limitations that contribute to variability in findings and compromise replicability (1, 6, 13). These discrepancies primarily arise from experimental designs lacking rigorous comparisons between resting-state or task-related brain activity during hypnosis and equivalent awake control conditions, thereby hindering the dissociation of induction-specific and suggestion-specific neurophysiological patterns (11, 14). Although findings remain mixed, a broad consensus suggests that hypnosis engages frontal networks linked to top-down cognitive control, most consistently evidenced by increased theta-band spectral power (4–7 Hz) following induction (6, 15). However, rather than oscillatory frequency *per se* being central to hypnotic responsiveness, electroencephalographic (EEG) studies indicate that hypnosis modulates neural connectivity across multiple frequency bands, from delta (1–4 Hz) to gamma (>30 Hz), emphasizing connectivity patterns as a key mechanism (15, 16). Accordingly, studies using interregional coherence measures, such as the imaginary component of coherency (iCOH; 17), have linked distinct connectivity changes to hypnosis induction (18). For instance, Jamieson and Burgess (19) found that highly suggestible individuals exhibit increased theta connectivity in a central-parietal hub and reduced beta connectivity in fronto-central and occipital regions. More recently, Niedernhuber et al. (20) reported reduced interhemispheric frontoparietal connectivity in hypnosis experts, facilitating information transfer between the left parietal and right frontal regions, independently of hypnosis depth. These findings may suggest that hypnosis induction suggestions promote cortical information flow along a frontal-to-posterior hierarchy and enhances frontoparietal integration, supporting attentional modulation and immersive hypnotic experiences, though replication is needed.

Variability in mental strategies when responding to specific hypnotic suggestions further poses a significant empirical issue, as similar behavioral outcomes may arise from different underlying cognitive and neural processes (7, 21, 22). This issue is exemplified in challenge-ideomotor paradigms, where participants are asked to imagine their arm becoming stiff like a bar of iron, and then attempt to bend it, creating a conflict between actual somatosensory feedback (the arm can bend) and the simulated perception of rigidity (a bar of iron cannot bend) (9, 23). Behavioral data from a single suggested-trial (try to bend the iron bar) revealed two distinct motor catalepsy strategies: one group exhibited arm trembling, while another showed no tremors, despite both reporting similar experiential responsiveness and an altered sense of agency (23, 24). This variability highlights the need for neurophenomenological approaches that integrate first-person subjective experiences with neural data (25), offering a relevant framework to disentangle the mechanisms underlying ideomotor phenomena (12, 26). To date, neuroimaging studies using functional magnetic resonance imaging have predominantly examined arm-paralysis ideomotor suggestions (without the challenged to bend it), comparing them to voluntary simulated motor inhibition to elucidate the neural mechanisms underlying altered motor control in hypnosis and the experience of involuntariness (27-30). Briefly, the results indicate that hypnosis enhances prefrontal activity linked to executive control, while suggested paralysis involves increased precuneus activation and reduced motor–premotor connectivity, supporting a role for self-monitoring over direct motor inhibition (27, 29, 31). As neural changes in hypnosis are primarily shaped by connectivity patterns, further EEG research is essential to deepening our understanding of ideomotor suggestion responsiveness, with the challenged paradigm offering a framework to disentangle the underlying mental strategies.

Overall, the neurophysiological and subjective changes underlying the top-down control that governs responsiveness to induction and ideomotor suggestions remains poorly understood. This study aimed to address this gap through two main objectives. First, we examined the neurophysiological dynamics underlying induction-type suggestions, focusing on the transition from light to deepening hypnotic states, compared to resting-state. Second, we investigated the neurophenomenological underpinnings of challenge-ideomotor suggestions, specifically arm-induced hypnotic catalepsy, relative to a simulated awake condition. To address these research questions, we employed a within-subject controlled randomization design and a multimodal assessment encompassing neural, physiological, behavioral, and first-person subjective measures.

## RESULTS AND DISCUSSION

### Induction state: Theta-coupled fronto-parietal network reorganization tracks hypnotic depth and vagal disengagement

EEG spectrum analysis with pairwise comparison between the Rest, Light and Deep states, using a cluster-based permutation test, revealed a decrease in alpha power between the Rest and Light states over electrodes mainly located in the parieto-occipital regions: PO3, POz, PO4, PO8, P6, P2, CPz, and CP4 (1 < *t*-values < 5, *P* = 0.041; Fig. 2*A*). Then, a generalized theta power enhancement was observed during the deepening state across all electrodes (25 < *t*-values < 175, *P* = 0.001; Fig. 2*B*).

**Fig. 1.**
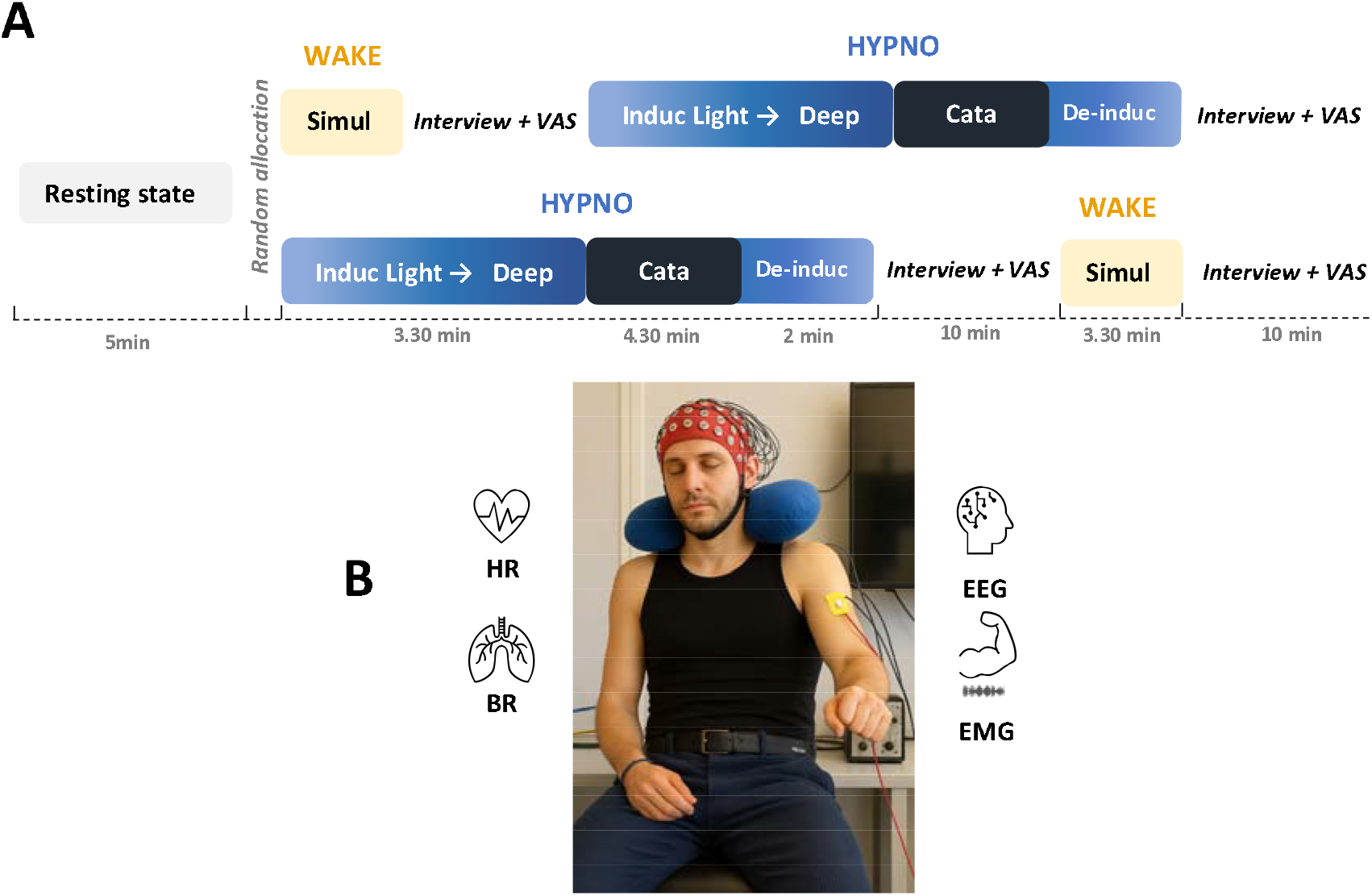
Experimental design and multimodal neurophysiological assessments. **A**. Following resting-state (Rest), participants were randomly assigned to two counterbalanced experimental conditions, with the WAKE session involving the awake simulation of impossible arm-bending trials (Simul), and the HYPNO session consisting of a hypnotic induction included a light induction followed by a deepening state using neutral induction-type suggestions and an ideomotor suggestion state targeting the left-arm cataleptic trials (Cata). Each session concluded with a first-person phenomenological interview and visual analog scale (VAS) to collect subjective experiences. **B**. Neurophysiological and multimodal assessments included continuous recordings of electroencephalography (EEG), heart rate (HR), breathing rate (BR), and electromyographic (EMG) activity of the biceps, and triceps muscles. Note: the face shown has been replaced with an AI-generated image for anonymization purposes.

**Figure 2.**
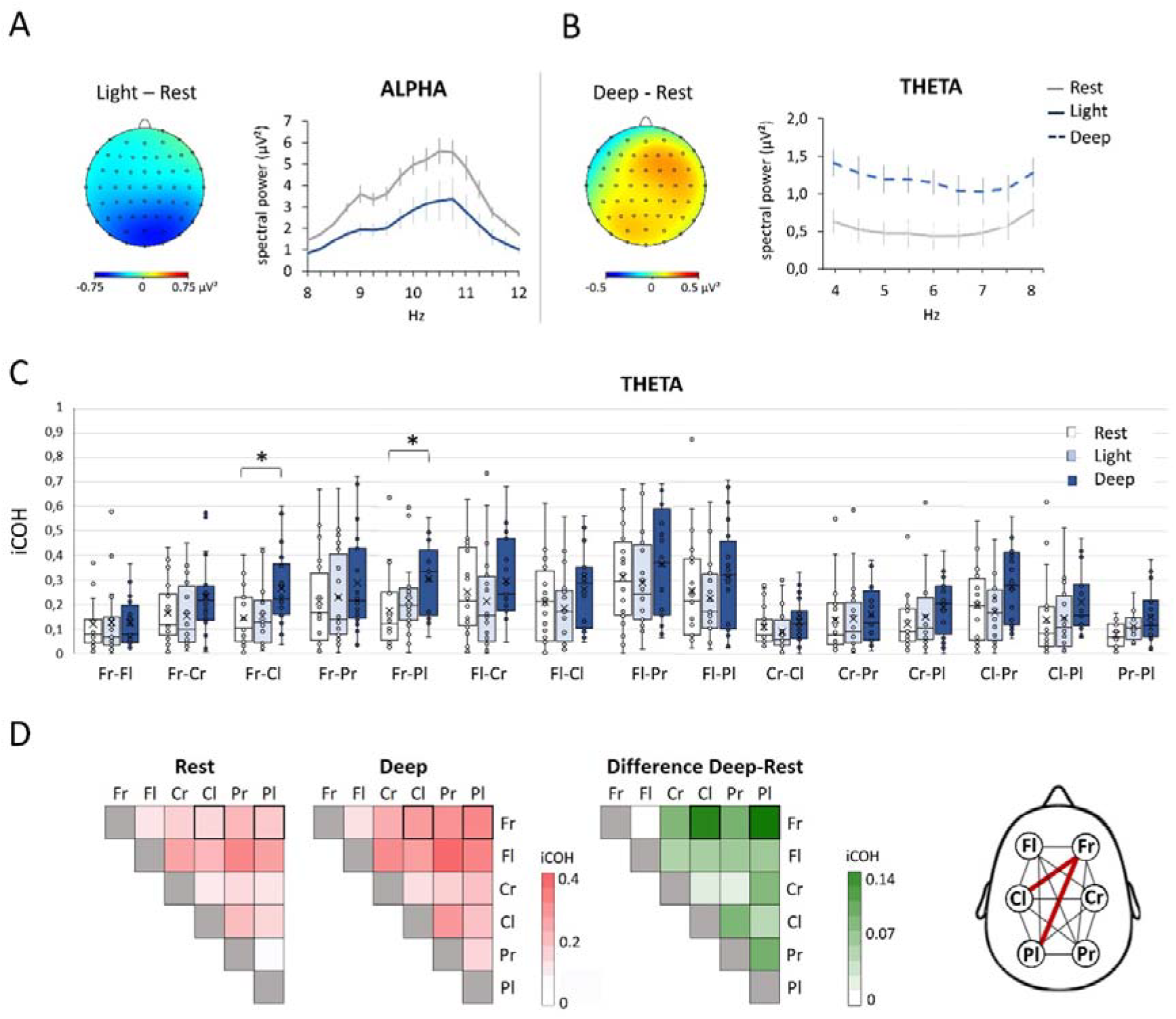
Spectral power and iCOH changes between resting state and light to deep hypnosis induction. **A**. Spectral power analysis across all electrodes, showing topographic and graphical differences in alpha power between the Light and Rest states, and **B**. theta power differences between the Deep and Rest states. **C**. Theta-band iCOH changes across Rest, Light, and Deep states for each brain region pair, with boxplots showing iCOH distribution, with the cross indicating the mean and the line the median. **D**. Heatmaps display iCOH values for Rest and Deep states on a red gradient, while the green map highlights differences between the two states, with significant changes in bold. Brain connectivity map illustrates the increase in connectivity in the theta band, during the Deep compared to Rest state between right frontal region (Fr) and left central and parietal regions (Cl and Pl respectively). Fl : right frontal; Cr : right central; Pr : right parietal regions. * *P* < 0.05.

To test for connectivity differences, imaginary coherence (iCOH) values were compared between the Rest, Light and Deep states, across pairs of regions (left and right frontal : Fr, Fl, central : Cr, Cl, and parietal : Pr, Pl regions). In the theta frequency band, the iCOH revealed a main effect of states (*F*_(2,38)_ = 4.145, *P* = 0.024), of regions (*F*_(14,266)_ = 11.822, *P* < 0.001), and a states*regions interaction (*F*_(28,532)_ = 1.507, *P* < 0.031; Fig. 2*C*). Post-hoc comparisons revealed that, during the deepening state, the right frontal region exhibited significant greater connectivity with the left parietal (*P* = 0.016) and left central regions (*P* = 0.034) in the theta band, compared to Rest (Fig. 2*D*). For the delta, alpha, beta and gamma frequency bands, only a main effect of the regions was observed (δ: *F*_(14,266)_ = 8.412, *P* < 0.001; α: *F*_(14,266)_ = 10.243, *P* < 0.001; β: *F*_(14,266)_ = 5.245, *P* < 0.001; γ: *F*_(14,266)_ = 4.200, *P* < 0.001).

Our findings showed that during the transition from *Rest* to *Light* hypnosis, we observed a decrease in alpha power over parieto-occipital regions, consistent with reduced processing of external stimuli and a reallocation of attentional resources toward internally guided states (20, 32). This early modulation likely marks the onset of perceptual decoupling and attentional redirection, preparing the neural landscape for both the absorption of suggestion-based content and the dissociation from the external environment (19), a process that exhibits neurophysiological similarities to the hypnagogic transition at sleep initiation (33, 34). Notably, these neural changes occurred despite an increase in breathing rate and no modulation of cardiac vagal tone as compared to Resting state (Supplementary data and Fig.1). The relative stability of these physiological markers during the *Light* state suggests that this initial hypnotic state does not reflect passive relaxation, but rather a cognitively engaged condition characterized by reoriented attentional control without parasympathetic activation. This view contrasts with traditional models framing hypnosis as a low-arousal state and aligns with recent neurocognitive perspectives emphasizing active top-down modulation during early stages of induction (1). However, the modulation of alpha activity in hypnosis remains debated as highlighted in the recent De Pascalis’ review (16), which emphasizes inconsistent findings across studies and suggests that such variability may, in part, reflect individual differences in hypnotizability (35).

As hypnosis deepens, a broader transformation emerges. The *Deep* state is characterized by a robust increase in theta power across the scalp, a frequency consistently associated with sustained attention, mental imagery, and cognitive flexibility (16). This spectral enhancement was accompanied by increased theta-band functional connectivity between the right frontal region and left parietal and central areas, mirroring the frontoparietal network organization identified in neuroimaging studies as critical for hypnotic absorption and altered self-awareness (2, 20, 36). Such cross-hemispheric coupling suggests the recruitment of large-scale control networks supporting internal representation and executive monitoring. Crucially, this transition to a more internally synchronized state was accompanied by increased vagal modulation, as reflected by a reduction in heart rate variability. This finding contrasts with the parasympathetic enhancement typically observed in meditation or relaxation-based hypnosis (36, 37), and suggests that hypnotic depth does not rely on autonomic quiescence. Rather, the observed pattern indicates a functional dissociation: enhanced neural coordination for cognitive control and mental imagery, paired with reduced parasympathetic tone which possibly reflects the sustained mental effort required to maintain immersive, suggestion-driven mental states.

Taken together, these findings suggest that hypnotic induction is not a unitary relaxation response but a staged brain–body transition. It begins with selective sensory decoupling and evolves into a state of large-scale network integration and attentional absorption, orchestrated through verbal suggestions. This process engages specific oscillatory and connectivity mechanisms while maintaining and even reducing parasympathetic tone, highlighting hypnosis as a distinctive neurocognitive state with functional autonomy from classic relaxation paradigms (38).

### Cataleptic state: Behavioral and phenomenology responsiveness during challenged ideomotor suggestions

In the Cata state, 20 out of 23 participants successfully responded to the ideomotor suggestion of ‘*not being able to bend the iron bar*’ across all 10 trials, with a subjective rating of cataleptic phenomenon of 9.2 ± 1.5 out of 10 for the left arm and 2.4 ± 0.7 for the right arm. The level of hypnotic depth in the HYPNO condition was 7.43 ± 1.18 out of 10. As expected, most were classified as *Tremblers* (n = 16), exhibiting muscle tremors during the Cata trials, while a smaller subset, the *Non-Tremblers* (n = 4), showed no visible tremors. This online observation of trembling behavior was supported by post hoc surface EMG analysis and participants’ subjective reports regarding their motor strategies. First, Root Mean Square (RMS) values, which quantify muscle activity intensity by measuring the overall amplitude of the EMG signal, were compared across Simul and Cata trials. No main effects were found for muscle (*F*_(1,35)_ = 1.329, *P* = 0.257), states (*F*(1,35) = 0.847, *P* = 0.364), or their interaction (*F*_(1,35)_ = 2.238, *P* = 0.144). To assess dynamic changes in muscle tone, a coefficient of variation of RMS values was computed for each participant. This analysis revealed greater RMS variability in *Tremblers* compared to *Non-Tremblers* for both the biceps (*U* = 5.05, *P* < 0.001) and triceps brachii (*U* = 10.00, *P* = 0.049). These results suggest that *Tremblers* alternated between contraction during trials and partial relaxation between trials, whereas *Non-Tremblers* maintained sustained muscle contraction throughout.

Further insights were obtained from a first-person interview conducted after both the HYPNO and WAKE conditions focused on mental strategies used during the Cata and Simul trials. The post hoc analysis revealed two main findings, based on a coding scheme developed following a grounded theory approach (39), which identified recurring themes directly from participants’ first-person accounts: (1) the mental strategies employed during the Cata and Simul conditions were inconsistent both within and across participants; and (2) *Tremblers* and *Non-Tremblers* differed in their experiential reports during Cata trials. Regarding the main thematic clusters, *Tremblers* frequently described attempting the movement despite perceiving it as involuntary or constrained, whereas *Non-Tremblers* typically reported not trying at all, either because it felt impossible from the outset or due to an inability to initiate the motor command.

These differences in mental strategies were also reflected in the subjective rating of perceived strength exertion. A paired t-test conducted across all participants comparing the subjective ratings between the Cata and Simul conditions revealed no significant difference (t = −0.95, *P* = 0.36). However, when examining individual responses, it was observed that four participants, identified as Non-Tremblers, showed reversed ratings compared to the majority, reporting lower perceived strength exertion during Cata compared to Simul.

Our findings showed that during the Cata trials, *Tremblers* and *Non-Tremblers* exhibited distinct motor responses, with the former showing rhythmic muscle oscillations both during and between trials, and the latter maintaining sustained muscular rigidity throughout. These patterns are consistent with previous studies reporting similar muscular distinctions, although those were limited to single-trial ideomotor suggestion challenges (9, 23). The present findings extend previous observations by revealing these motor distinctions across multiple trials and, through first-person interviews, further suggest that they are associated with distinct mental strategies. Importantly, all participants were able to recall and describe their experience during the hypnotic Cata trials, consistently reporting altered sense of agency, which was accompanied either by an effective motor attempt (*Tremblers*) or by the absence of any movement (*Non-Tremblers*). These findings are consistent with Deeley et al. (29), who demonstrated preserved motor intention, as *Tremblers* here, alongside involuntary inhibition during suggested limb paralysis. Similarly, in our study, all participants retained awareness of the hypnotic experience and reported involuntariness. However, our behavioral distinction between *Tremblers* and *Non-Tremblers* extends this model by revealing a gradient in motor output and sense of agency. While both groups shared a dissociation from volitional control, only *Non-Tremblers* exhibited a complete absence of movement, suggesting a deeper disruption in the sense of agency and motor ownership. This divergence indicates that awareness of the experience may remain intact, whereas the phenomenological access to agency can vary across individuals, even under the same suggestion.

### Challenged ideomotor suggestions reduced delta and enhanced gamma connectivity in right fronto-parietal networks

Given the differences in mental strategies used during Cata trials, only *Tremblers* were included in the following EEG analyses (n = 16). Power spectrum differences between the Cata and Simul conditions did not reveal any significant clusters for any frequency bands. Changes in iCOH in the delta and gamma band revealed a main effect of states (δ: *F*_(1,15)_ = 5.843, *P* = 0.03; γ : *F*_(1,15)_ = 5.08, *P* = 0.04) and a main effect of regions (δ: *F*_(14,210)_=6.719, *P* < 0.001; γ: *F*_(14,210)_ = 3.26, *P* < 0.001), but no states*regions interaction (δ: *F*_(14,210)_ = 1.295, *P* = 0.21; γ: *F*_(14,210)_ = 1.163, *P* = 0.31). Results showed that during Cata, the right frontal region exhibited lower connectivity with the right parietal (*P* = 0.023) and central (*P* = 0.033) regions in the delta band compared to Simul (Fig. 3*A*). Conversely, in the gamma band, the right parietal region showed increased connectivity with the right frontal (*P* = 0.036), right central (*P =* 0.031), and left parietal (*P* = 0.033) regions during Cata compared to Simul (Fig. 3*B*). For the theta, alpha, and beta frequency bands, only a main effect of regions was observed (θ: *F*_(14,210)_ = 3.67, *P* < 0.001; α: *F*_(14,210)_ = 8.33, *P* < 0.001; β: *F*_(14,210)_ = 5.28, *P* < 0.001).

**Figure 3:**
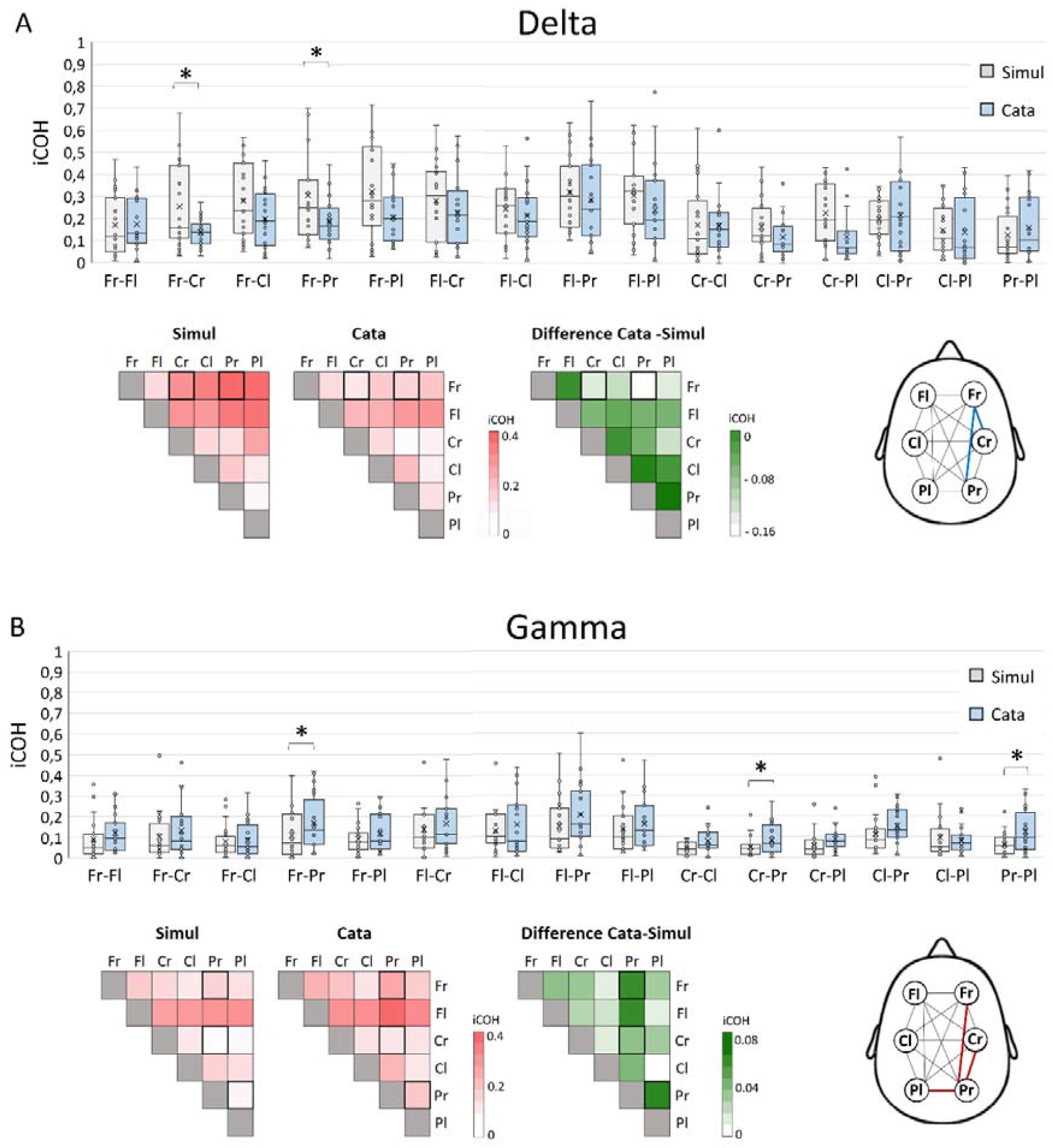
Differences in imaginary coherence (iCOH) between Simulation and Catalepsy trials for delta and gamma frequency bands across brain regions. **A**. Boxplots representing iCOH distributions across subjects, with the central cross indicating the mean and the line representing the median. Heatmaps display iCOH values for Simul and Cata states on a red gradient, while the green map highlights differences between the two states, with significant changes in bold. Brain connectivity map illustrates lower delta-band iCOH in Cata compared to Simul (blue) between right frontal (Fr) and right central (Cr) and parietal (Pr) regions. **B**. Similar representations for the gamma frequency band: Boxplots show iCOH distributions, with the mean and median indicated. Heatmaps illustrate iCOH for Simu and Cata on a red gradient, with a green map representing the state differences and significant effects highlighted in bold. Brain connectivity map illustrates the increase in connectivity in the gamma band, during Cata compared to Simul between right parietal region (Pr) and right central (Cr), right frontal (Fr) and left parietal (Pl) regions. \**P* < 0.05.

Findings revealed that Tremblers exhibited a peculiar reorganization of cortical connectivity, marked by increased gamma-band coupling between the right frontal, right parietal, and left parietal regions, and a concurrent decrease in delta-band connectivity between right frontal and right parietal sites. This pattern may provide support the theta-gamma oscillation model of hypnosis proposed by Jensen et al. (15), which posits that slow oscillations can modulate faster dynamics through phase-coupling mechanisms. While their framework primarily highlights theta–gamma coupling, our findings extend this model by indicating that delta–gamma connectivity may also play a role in the functional integration of sensorimotor networks during ideomotor challenge suggestions.

From a mechanistic perspective, these patterns may reflect a weakening of low-frequency coordination in favor of localized gamma synchrony, which facilitates sensorimotor binding and executive monitoring during volitional conflict under ideomotor challenged conditions (40, 41). Notably, Jensen et al. (15) describe slow-wave activity as supporting the reactivation of declarative memory, including somatosensory representations, which could be mobilized here to reinstate the sensations such as “arm rigidity”. Gamma oscillations, in turn, have been associated with the binding of sensorimotor representations and the integration of conscious experience (42, 43), and under conditions of high cognitive demand or sensorimotor conflict in high susceptible hypnotic individual (18). In this context, delta suppression may reflect a release from passive low-frequency coordination, freeing cortical resources for gamma-mediated top-down monitoring and integration of action-related representations during conflict resolution (16). Additional support for this account comes from the observed reduction in heart rate variability during Cata relative to Simul, consistent with heightened cognitive demand. This interpretation aligns with prior neuroimaging studies of hypnotic motor inhibition, which have shown that attempts to act under suggestion engage prefrontal executive networks⍰(29, 44), potentially overriding low-frequency inhibitory rhythms such as delta. Notably, the observed oscillatory dissociation was localized in right fronto-parietal regions contralateral to the suggested left arm, corresponding to areas known to support self-monitoring, motor inhibition, and altered experiences of agency during hypnosis (44). Thus, the emergence of enhanced gamma coupling within right-hemisphere fronto-parietal circuits, contralateral to the left arm targeted by the challenged ideomotor suggestions, could reflect the active attempt to reconcile prediction errors in motor agency by sustaining a coherent sensorimotor representation aligned with the ideomotor suggestions.

Our observed oscillatory dynamics, behavioral responses, and subjective reports lend further support to recent predictive coding frameworks of hypnotic ideomotor suggestion (13, 45). Although both Tremblers and Non-Tremblers reported a loss in the sense of agency, the two groups diverged markedly in their behavioral and phenomenological profiles. Tremblers attempted to perform the suggested movement despite its perceived impossibility, producing rhythmic motor activity, while Non-Tremblers exhibited no overt movement and described a sense of motor command inhibition or blockage. From a predictive coding perspective, this dissociation reflects alternative strategies for minimizing sensorimotor prediction error. Tremblers may engage in active inference, generating proprioceptive prediction errors through motor attempts in an effort to reconcile internal models with incoming sensory feedback (46). This suggests a lower precision weighting of the “non-agency” prior induced by the suggestion, permitting partial expression of bottom-up motor signals. In contrast, Non-Tremblers appear to rely on perceptual inference, assigning greater precision to the suggestion-based prior and attenuating the influence of contradictory sensory feedback. This contrast exemplifies the core mechanism of precision weighting in predictive coding, wherein the brain dynamically balances top-down expectations and bottom-up evidence to maintain coherence in motor and experiential representations (47).

Our findings reveal that hypnotic suggestions modulate brain–body dynamics through frequency-specific network reorganizations and divergent sensorimotor strategies. This supports predictive coding models of agency and underscores hypnosis as an active, non-unitary neurocognitive state. By integrating neurophysiological and phenomenological data, this study offers novel insights into how conscious experience is shaped by top-down processes. These results not only refine the theoretical understanding of hypnosis and volition, but also inform the development of individualized hypnotic interventions, particularly in contexts such as pain modulation, motor rehabilitation, and functional neurological disorders, where ideomotor suggestions may engage therapeutic reorganization of bodily awareness.

## MATERIEL AND METHOD

### Participants

An a priori power analysis indicated that a sample of 18 participants would be sufficient to detect a medium effect size in EEG connectivity (f⍰=⍰0.25) across repeated measures (i.e., state comparisons: Rest, Light, Deep), with α set at 0.05 and power (1⍰−⍰β) at 0.80. This estimate was based on effect sizes reported by Jamieson and Burgess (19) for theta and beta-band iCOH following hypnotic induction in a similar within-subjects design. To account for potential data loss due to EEG artifacts or protocol-related variability, we aimed for a sample of 23 participants.

The study protocol was approved by the Research Ethic committee of the University of Lyon (2023-10-19-002) and followed the principles expressed in the Declaration of Helsinki. Consent forms for participation in initial hypnotizability screening and experimental session were provided. Subjects were excluded if they reported any history of neurological or psychiatry disease, or took medication or drugs. We screened 40 healthy right-handed individuals using the Harvard Group Scale of Hypnotic Susceptibility: FormA (HGSHSA) [4]. This test involved a structured hypnotic induction followed by a series of 12 suggestions targeting sensory, motor, and volitional alterations. Only participants who achieved a minimum score of 6 out of 12 items, including successful compliance with the challenged ideomotor suggestion, were included in the experimental study. Our final sample was composed by 23 healthy right-handed (11 women, mean age of 24.75 ± 3.78) with medium to high level of hypnotizability (8.80 ± 1.82). Three participants were excluded from the analyses as the ideomotor challenge suggestion failed during the experiment, leading them to bend their arm during the Cata trials.

### Experimental Design

The experiment was conducted in a quiet, controlled environment to minimize external disturbances. One experimenter (JG) was responsible for ensuring the real-time acquisition and monitoring of multimodal assessments, while a second experimenter (UD), a trained hypnotist, administered the hypnotic instructions and guided the participants throughout the experimental session including first person interview. This division of roles ensured the seamless integration of data collection and the consistent delivery of hypnotic protocols. The participant was seated in an armchair, a cushion placed around the neck, with continuous recordings of heart rate and breathing rate (HR/BR), left-arm electromyography (EMG), and 64-channel EEG throughout the session. The experiment began with a five-minute eyes-closed resting state, during which participants were instructed to remain still and avoid focused thoughts, providing a baseline for neurophysiological measures, including EEG and HR/BR. Participants then proceeded with the waking ordinary state of consciousness condition (WAKE), followed by the hypnotic condition (HYPNO), with the order of conditions counterbalanced across participants. Before each condition, participants assessed their current state of alertness via the Stanford Sleepiness Scale (SSS, 48); with a minimum level at 3 required (i.e. relaxed; awake; not at full alertness; responsive).

During the WAKE condition, the experimenter showed the motion of bending the left forearm over the upper arm and instructed participants to simulate being unable to perform this movement, acting “*as if they were trying to bend their left arm but were unable to do so*,” while emphasizing that they could *“feel free to perform the simulation in their own manner*.” Then, participants were asked to raise their left arm, stretched out in front of them, with their fist and eyes closed. Each simulated trial (Simul) began with a verbal ‘go,’ lasted approximatively four seconds (± 1 s), and was followed by a five-second break during which subject maintained the position. A total of ten trials were conducted, and condition lasted 3.28 ± 0.49 minutes.

The HYPNO condition began with induction suggestions based on Gruzelier’s model (49), starting with visual fixation. Participants were instructed to extend their left arm forward and focus their gaze on their thumb until the arm naturally lowered, their eyelids gradually closed, and their hand rested on their thigh. On average, the duration of attentional induction was 1.30 ± 1.02 minutes. Then, the relaxation and multimodal imagery induction-type suggestions used to guide the transition from light to deepening hypnosis (see the full script in Supp file) beginning with *‘You can focus on your exhalation… Every time you exhale, pay attention to how your muscles relax*… *Continue to let this sensation of relaxation spread…*’. The deepening state was initiated with a counting sequence like in the standardized Haward test, inviting the participant to move into an even deeper state of comfort with each number beginning from 20. During this phase, the suggestions aimed to increase the participant’s internal focus absorption, and the dissociation of consciousness between the physical and the mental like : *‘You will move into a state of comfort in which you will be able to perform movements I will ask you to make…Pay only attention to my voice…In this state of deep relaxation, you may find yourself feeling both present and elsewhere… both focused and distracted … neither tense nor relaxed…*’. On average, the Light to Deep induction state was 3.23 ± 0.24 minutes.

Then levitation suggestions were provided for the left arm and hand, and once the desired position in front of the participant was achieved, the experimenter began ideomotor suggestions, indicating that their left arm was becoming so rigid, like a ‘metal bar’ that it could no longer be bent (see the script in supplementary file). They were then invited to attempt bending the ‘metal bar,’ which was suggested as impossible, for four seconds (± 1 s) during ten cataleptic trials (Cata), and rigidity suggestions were maintained across the 5-s inter-trials to keep the adherence of the subject. For instance: *‘It [subject’s arm] is becoming stiff* … *more and more stiff* … *rigid* …*like a bar of iron* … *and you know how difficult how impossible it is to bend a bar of iron like your arm… Try to bend it [4 s]… you can stop trying’*. The termination of the hypnosis followed a standardized procedure, including a suggestion to shift focus back to the physical environment and restore typical expectations regarding the influence of one’s actions and perception of external stimuli. The duration of challenged ideo-motor condition was 4.36 ± 0.57 minutes.

After both the WAKE and HYPNO conditions, a structured debriefing session was conducted to gather first-person phenomenological data on participants’ subjective experiences and motor control strategies. This involved targeted questions such as *“What/how do you do when you hear ‘go’?”* or *“What/how do you do when you hear ‘try to bend the bar’?”*, depending on the condition just completed. The interview was conducted by UD, who is trained in phenomenological interviewing techniques aimed at guiding participants to explicit their subjective experiences, focusing on the action (mental/motor) and describe them in rich detail (50) and intentionality (self-agency). The primary objective of this phenomenological approach was to examine the cognitive strategies underpinning motor control mechanisms and to explore the intentionality i.e. sense of agency, with the entire debriefing lasting approximately five minutes. They were further asked to rate on a VAS their left arm muscular fatigue, and their subjective strength deployed during trials. Participants were also asked to rate the subjective depth of their hypnotic experience following the Hypno condition, using a 0–10 scale, where 0 indicated “not hypnotized at all” and 10 corresponded to “as deeply hypnotized as you have ever been”. The time between the two conditions was 9.04 ± 0.52 minutes. Finally, to confirm the effectiveness of the arm rigidity suggestion, participants completed the Catalepsy Questionnaire (51) assessing their perception of both arms during the hypnotic suggestion. The closer the score was to 10, the more effective the arm rigidity suggestion was in the affected arm. Two isometric maximum voluntary contractions of the triceps and biceps brachii were performed, with 2 min of recovery between each, to allow the normalization of the EMG data for analysis.

### EEG recordings and preprocessing

EEG was recorded from 64 EEG electrodes (ActiCAP snap, BrainProducts®, Munich, Germany) localized according to the 10–20 system on participants’ heads. The impedance of each electrode was kept below 10 kOhm. The initial sampling rate was 1000 Hz, and data were down-sampled off-line to 256 Hz. Reference was on mastoids (TP9 and TP10) and converted to average reference offline. The EEG signal was amplified by BrainAmp amplifiers and recorded with the BrainVision Recorder software (Brain Products®, Munich, Germany). The EEG data were processed and analyzed offline using the BrainVision® System (BrainProducts®, Munich, Germany). A 1 Hz high-pass filter and a 50 Hz notch filter were applied to the EEG signal. EEG recordings for each condition were segmented into 3-s epochs. Independent component analysis was used to remove eye movements and muscles artifacts. Any segments containing artifacts over 250 μV were discarded from the analysis. Fast Fourier Transforms (FFT) and phase-coherence analysis were performed on the artifact-cleaned epochs of the Rest, Light, Deep states, and Cata and Simul conditions.

### Spectral analysis

Each 3-s EEG segment was first windowed with a Hanning tapering window prior to computing the power spectra using the FFT in five spectral band: δ (0.5-3.5), θ (4-7.5 Hz), α (8-12 Hz), β (13-30Hz) and γ (31-80 Hz) and computed for each electrode. The power content, expressed as μV^2^ for each condition, was determined as the average power across the 3-s segments of the EEG. For analysis, the relative power was calculated by dividing the sum of spectral power from all bins for each frequency bin within 0 and 80 Hz and each electrode.

### Functional connectivity analysis

Functional connectivity was studied using EEG imaginary phase-coherence analysis between six pairs of regions (Frontal left : mean of FP1, AF3, AF7, F3, F5, F7; Frontal right : mean of FP2, AF4, AF8, F4, F6, F8; Central left : mean of FC3, FC5, FT7, C3, C5, T7, CP3, CP5, TP7; Central right : mean of FC5, FT7, C3, C5, T7, CP3, CP5, TP7; Parietal left : mean of P3, P5, P7, PO3, PO7,; Parietal right : mean of P4, P6, P8, PO4, PO8) for all conditions (Rest, Light, Deep, Cata and Simul) and for each frequency bands (δ, θ, α, β and γ). Coherence is defined as the square of the cross-spectrum between electrodes, divided by the product of the power spectra of the individual electrodes (52). This calculation provides a measure of the consistency of the phase relationship between two signals, with values ranging from 0 to 1. However, the measure of coherence is very sensitive to volume conduction, which can lead to erroneous correlation estimates. To address this issue, we focused on imaginary coherence (iCOH), which has been proposed as an effective method for assessing brain connectivity based on the frequencies of brain signals (52). The imaginary component of coherency is obtained by taking the imaginary part of the cross-spectrum of the electrodes and dividing it by the square root of the product of the individual electrodes’ power spectra. Thus, after performing the FFT of the signal for each spectral band power, the absolute value of the imaginary result was computed across the 3-s segments of the EEG. The total iCOH was calculated separately for each frequency band.

### Heart rate variability and breathing rate measures

The Hexoskin Biometric Shirt (Carré Technologies Inc., Montreal, Québec, Canada) was used with the Hexoskin Smart Device (datalogger) to measure the participants’ average heart rate (HR) and breathing rate (BR). This wearable monitor is a compression shirt with built-in cardiac and breathing sensors sit around the thorax and abdomen. The cardiac sensors are analog ECG recording at 256 Hz and the respiratory sensors are dual channel respiratory inductance plethysmography sensors recording at 128 Hz. HR was detected from the ECG and recorded every second in units of beats per minute. The BR was recorded every second in units of breaths per minutes (bpm). The datalogger collects data from the sensors and stores it internally. Markers were added at the beginning and end of each state. At the end of the experimental session, the data was extracted from the datalogger to a computer via USB. HR analysis, including artifact detection and extraction of time and frequency domains, was performed using MATLAB software (R2019a®). In the time domain, the root means square successive difference (RMSSD, in ms) was calculated. In the frequency domain, the power in the low (LF: 0.04-0.15 Hz) and high (HF: 0.15-0.40 Hz) frequency bands were quantified, and the LF/HF ratio was subsequently calculated.

### Surface electromyography

Five surface EMG electrodes (Asept foam adhesive electrodes, medico electrodes international LTD) were attached to the subject’s left arm over the biceps brachii and long head of the triceps brachii in accordance with Surface Electromyography for the Non Invasive Assessment of Muscles guidelines (53). The reference electrode was attached to the wrist bone. Prior to the application of the electrodes, the areas around the electrode attachment sites were shaved and rubbed so that skin impedance could be greatly reduced. A cotton pad soaked with alcohol was then used to clean the skin. The electrodes were placed parallel to the muscle fibers with inter-electrode distance of 2 cm. EMG signal was acquired using data acquisition software LabChart 8 and sampled at a frequency of 2 kHz. Then, the EMG signal was filtered by a 50 Hz notch filter to remove line noise and a 20–500 Hz fourth-order band-pass filter. From the EMG signal of the Cata and Simul conditions, the root mean square (RMS) was calculated over a 3-second period for each trial. A Maximum Voluntary Contraction-normalization was performed to allow comparison of RMS values between subjects. Thus, obtained RMS values represent the percentage of maximum force exerted by each participant during two isometric voluntary contraction per muscle. Average over all subjects (n = 20) was computed for the Simul and Cata conditions.

### Phenomenological analysis

To analyze participants’ experiential reports from the explicitation interviews conducted after the Cata and Simul trials, we used an emergent coding scheme following a grounded theory approach (Glaser, 1992). The goal of the interviews was to probe both *what* participants experienced during the trials and *how* they mentally and physically enacted the suggested tasks, with a specific focus on motor strategies and intentionality. Two raters, one of whom was blind to participants’ experience, analyzed the full transcript of all reports. Each rater initially reviewed the data independently and developed a list of non-overlapping experiential dimensions that could classify all responses with the minimal number of distinct categories. Dimensions were then compared and discussed, focusing specifically on motor engagement and intention during the trials. Inter-rater agreement was particularly strong for motor-related experiential dimensions, which were consistently and independently identified by both raters. Minor discrepancies on other categories were resolved through discussion.

The analysis yielded a clear distinction between two participant profiles in the Cata condition. Among the 16 Tremblers, most described attempting to bend the bar, even while acknowledging that doing so felt difficult or impossible. In contrast, the 4 Non-Tremblers typically reported not attempting the movement at all, often describing an experience in which the motor command could not be initiated or failed to reach the arm.

In the Simul condition, where participants were asked to simulate the same task in a wakeful, non-hypnotic context, all individuals reported trying to perform the movement voluntarily. Most described engaging in active blocking strategies—such as mentally halting the command at a specific joint (e.g., the elbow or wrist)—while a few reported choosing not to move without invoking an internal sense of inhibition.

### Statistical analyses

Data analyses were performed with JASP®software (version 0.16.1.0). Differences in the power spectrum across all electrodes were statistically assessed using a cluster-based permutation test comparing conditions pairwise. The t-statistic was calculated for each cluster, which was defined as two or more spatially contiguous electrodes where the t-statistics of power spectrum exceeded a chosen threshold of alpha level of p < 0.05 and α_cluster_ = 0.01. A total of 1,000 permutations were performed to establish a null distribution of the test statistic. To test for connectivity differences, ANOVA_RM_ were first conducted on iCOH values between the three states (Rest, Light and Deep) and then between Simul and Cata, across pairs of regions (Fr, Fl, Cr, Cl, Pr, Pl) for each frequency band separately. Two-way ANOVArm, with states (Cata vs Simul) as within factor and muscles (Biceps, Triceps) as between factor, was conducted on the RMS values from the EMG. To assess the difference in the coefficient of variation according to the trembling behavior, Mann-Whitney tests were computed for the Cata and Simul states, for both the biceps and triceps brachii.

To examine the effect of Rest, Light, and Deep states on one hand, and the Rest, Simul and Cata states on the other hand, on heart rate variability (i.e., RMSSD and LF/HF), and on BR, ANOVA_RM_ were performed.

For each ANOVAs, the Greenhouse–Geisser correction was applied when necessary. Post hoc analyses with Bonferroni correction for multiple comparisons were performed when significant effects or interactions were found following ANOVAs.

