## Supplemental data and figure 1 for "Decoding hypnotic consciousness: neural and experiential insights into induced and ideomotor suggestions"

Analysis of physiological BR across the induction revealed a main effect of phase (F_(1.879,31.939)_ = 3.862, *P* = 0.03), with a higher BR during the Light phase compared to Rest (*P* = 0.03). No differences were observed between Rest and Deep (*P* = 0.96), or between Light and Deep phases (*P* = 0.27; Supp Fig.1). Heart-rate variability analysis showed no effect of phase on the Low frequency/Hight frequency ratio (F_(2,21)_ = 0.76, *P* = 0.48). However, a main effect of the Root Mean Square of Successive Differences (RMSSD; F_(2,28)_ = 4.80, *P* = 0.016), with post-hoc analysis indicating lower RMSSD values during the Deep phase compared to Rest (Post-hoc test, *P* = 0.015), suggesting a reduction in parasympathetic activity during Deep phase. Noteworthy, breathing rate analysis revealed a main effect of phase (F_(1.837,31.222)_ = 6.253, *P* = 0.006), with a higher breathing rate during Cata compared to Rest (*P* = 0.005 ; Fig. 1), accompanied by a RMSSD reduction in Cata compared to Rest (F_(2.28)_ = 9.692, *P* < 0.001) and compared to Simul (*P* = 0.035), suggesting higher parasympathetic activity during Cata.

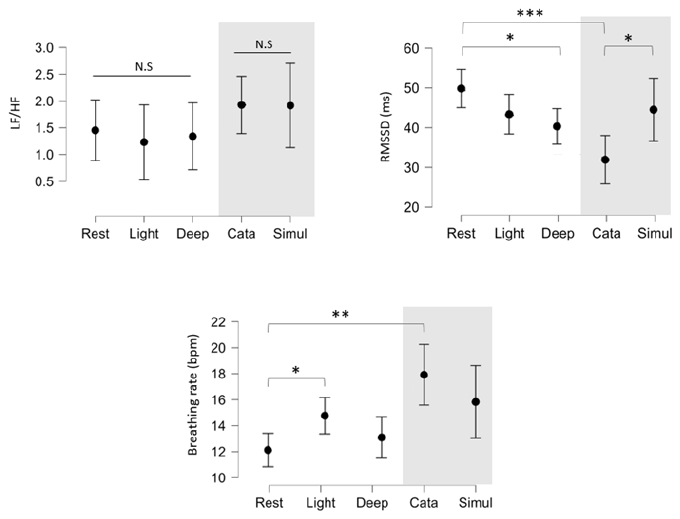

**Supp. Fig.1: Physiological responses across different phases of HYPNO condition and WAKE simulation**. A. Low frequency/High Frequency (LF/HF) ratio during Rest, Light, Deep, Catalepsy (Cata), and Simulation (Simul) phases. B. Root Mean Square of Successive Differences values (RMSSD) across phases, showing a significant decrease during Deep compared to Rest (*P < 0.05), and significant differences between Deep, Cata, and Simul (***P < 0.001, *P < 0.05). C. Breathing rate across phases, with a higher rate during the Light phase compared to Rest (*P < 0.05) and a global increase during Simulation (**P < 0.01). Data are presented as means ± SEM.
